# Dimeric form of SARS-CoV-2 polymerase

**DOI:** 10.1101/2021.03.23.436644

**Authors:** Florian A. Jochheim, Dimitry Tegunov, Hauke S. Hillen, Jana Schmitzova, Goran Kokic, Christian Dienemann, Patrick Cramer

## Abstract

The coronavirus SARS-CoV-2 uses an RNA-dependent RNA polymerase (RdRp) to replicate and transcribe its genome. Structures of the RdRp revealed a monomeric enzyme composed of the catalytic subunit nsp12, two copies of subunit nsp8, and one copy of subunit nsp7. Here we report an alternative, dimeric form of the coronavirus polymerase and resolve its structure at 4.8 Å resolution. In this structure, the two RdRps contain only one copy of nsp8 and dimerize via their nsp7 subunits to adopt an antiparallel arrangement. We speculate that the RdRp dimer facilitates template switching during production of sub-genomic RNAs for transcription.

---

Replication and transcription of the RNA genome of the coronavirus SARS-CoV-2 rely on the viral RNA-dependent RNA polymerase (RdRp)^1–5^. Following the structure of the RdRp of SARS-CoV-1^6^, structures of the RdRp of SARS-CoV-2 were obtained in free form^7^ and as a complex with bound RNA template-product duplex^8–11^. These structures revealed a monomeric RdRp with a subunit stoichiometry of one copy of the catalytic subunit nsp12^12^, two copies of accessory subunit nsp8^13^, and one copy of the accessory subunit nsp7^3,14^. Some studies additionally observed monomeric RdRp lacking one of the two nsp8 subunits and nsp7^6,9^. Here we show that the RdRp of SARS-CoV-2 can also adopt an alternative, dimeric form with two RdRps arranged in an antiparallel fashion.

To detect possible higher-order RdRp assemblies in our cryo-EM data for RdRp-RNA complexes, we wrote a script to systematically search for dimeric particles (Methods). We calculated nearest-neighbor (NN) distances and relative orientations of neighboring RdRp enzymes. In one of our published data sets (structure 3^11^), we detected many particles that showed RdRps with a preferred NN distance of 80 Å and a relative orientation of 180° (**Supplemental Figure 1a, b**), indicating the existence of a structurally defined RdRp dimer.

From a total of 78,787 dimeric particles we selected 27,473 particles that showed a strong RNA signal during 2D classification. The selected particles led to a 3D reconstruction at an overall resolution of 5.5 Å (**Supplemental Figure 1c**). We then fitted the obtained density with two RdRp-RNA complexes^10^, which revealed an antiparallel arrangement and RNA exiting to opposite sides of the dimeric particle. When we applied the two-fold symmetry during 3D reconstruction, the resolution increased to 4.8 Å (**Supplemental Figure 1d**).

The reconstruction unambiguously showed that both RdRps lacked one copy of nsp8, and thus each enzyme was comprised of only one copy of nsp12, nsp8, and nsp7 (**Supplemental Figure 2a**). The lacking nsp8 copy is the one that interacts with nsp7 in monomeric RdRp and had been called nsp8b^10^. The reconstruction showed poor density for the protruding second turn of the RNA duplex, the C-terminal helix of nsp7 (residues 63-73) and the sliding pole of the remaining nsp8 subunit (nsp8a, residues 6-110). These regions were mobile in both RdRp complexes and were removed from the model. Rigid body fitting of the known RdRp domains led to the final structure.

The structure of the antiparallel RdRp dimer showed that the two polymerases interact via their nsp7 subunits, with the nsp7 helices α1 and α3 (residues 2-20 and 44-62, respectively) contacting each other (**Figure 1**). Formation of the nsp7-nsp7 dimer interface is only possible upon dissociation of nsp8b, which liberates the dimerization region of nsp7 (**Supplemental Figure 2b**). The nsp7-nsp7 interaction differs from a previously described interaction in a nsp7-nsp8 hexadecamer structure^15^.

**Figure 1.**
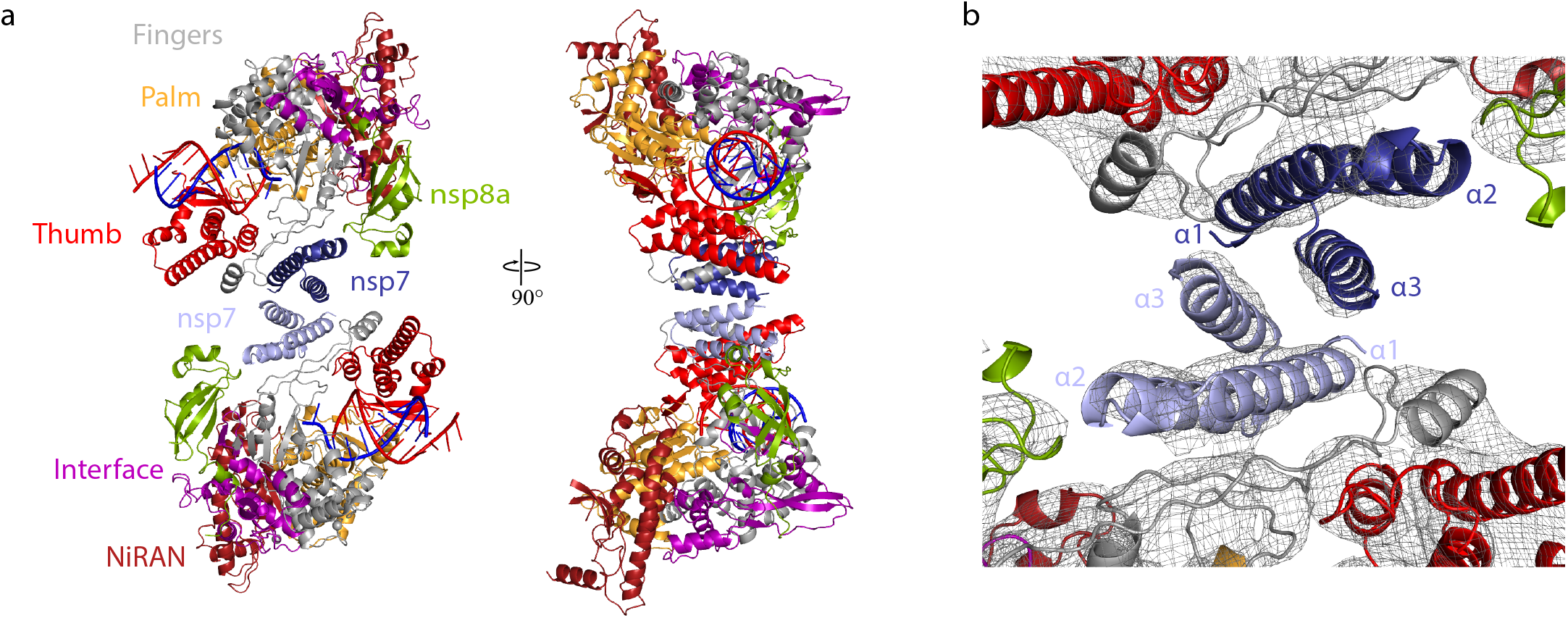
Structure of antiparallel RdRp dimer. **a** Two views of a ribbon model of the antiparallel RdRp-RNA dimer. Color code for nsp7, nsp8, nsp12 domains (NiRAN, interface, fingers, palm and thumb), RNA template (blue) and RNA product (red) is used throughout. Nsp7 subunits in the two RdRp monomers are colored slightly differently for the two monomers (dark and light blue, respectively). Views are related by a 90° rotation around the vertical axis. **b** Close-up view of nsp7-nsp7 dimerization interface. View is as in the left structure of panel a. The final cryo-EM density is shown as a black mesh.

The nsp7-nsp7 interaction is however consistent with formation of a (nsp7-nsp8)2 heterotetramer^16,17^ and its dissociation into a stable nsp7-nsp8 heterodimer that was suggested to bind nsp12^18^.

Frequently occurring mutations in the SARS-CoV-2 genome are predicted to influence formation of the RdRp dimer. The nsp12 mutation P323L^19^ coevolved with the globally dominating spike protein mutation D614G^20^, is found predominantly in severely affected patients^21^ and is predicted based on the structure to stabilize nsp12 association with nsp8a^20,22^. In contrast, the nsp7 mutation S25L is predicted to destabilize nsp7 binding to nsp8a^22^ and the nsp7 mutation L71F, which is associated with severe COVID-19^23^, may destabilize binding of the nsp7 C-terminal region to nsp8b. This suggests that mutations in RdRp subunits can influence the relative stabilities of the RdRp monomer and dimer.

Neither the catalytic sites nor the RNA duplexes are involved in RdRp dimer formation. It is therefore likely that the two RdRp enzymes remain functional within the dimer structure. The two RdRp enzymes in the dimer may thus be simultaneously involved in RNA-dependent processes. Unfortunately, we could not test whether the RdRp dimer is functional because we could not purify it.

We suggest that the RdRp dimer is involved in the production of sub-genomic RNA (sgRNA)^24–26^. In this intricate process, positive-strand genomic RNA (gRNA) is used as a template to synthesize a set of nested, negative-strand sgRNAs that are 5’ and 3’ coterminal with gRNA. The obtained sgRNAs are later used as templates to synthesize viral mRNAs. Production of sgRNAs involves a discontinuous step, a switch of the RdRp from an upstream to a downstream position on the gRNA template^27^. These positions contain transcription regulatory site (TRS) sequences^28–30^, but it is enigmatic how a single RdRp enzyme could ‘jump’ between these.

Our dimer structure suggests a model for sgRNA synthesis that extends a recent proposal^31,32^ (**Figure 2**). In the model, one RdRp of the dimer (RdRp 1) synthesizes sgRNA from the 3’ end of the gRNA template until it reaches a TRS in the template body (TRS-B). Due to the lack of one nsp8 subunit, the dimeric RdRp is predicted to have lower processivity than monomeric RdRp^10,14^ and this may facilitate TRS recognition. The viral helicase nsp13 could then cause backtracking of the RdRp^31,32^. Backtracking exposes the 3’-end of the nascent sgRNA, which is complementary to the TRS and may hybridize with another TRS located in the leader (TRS-L) at the 5’-end of the template. The resulting RNA duplex could bind to the second RdRp (RdRp 2) and sgRNA synthesis continues.

**Figure 2.**
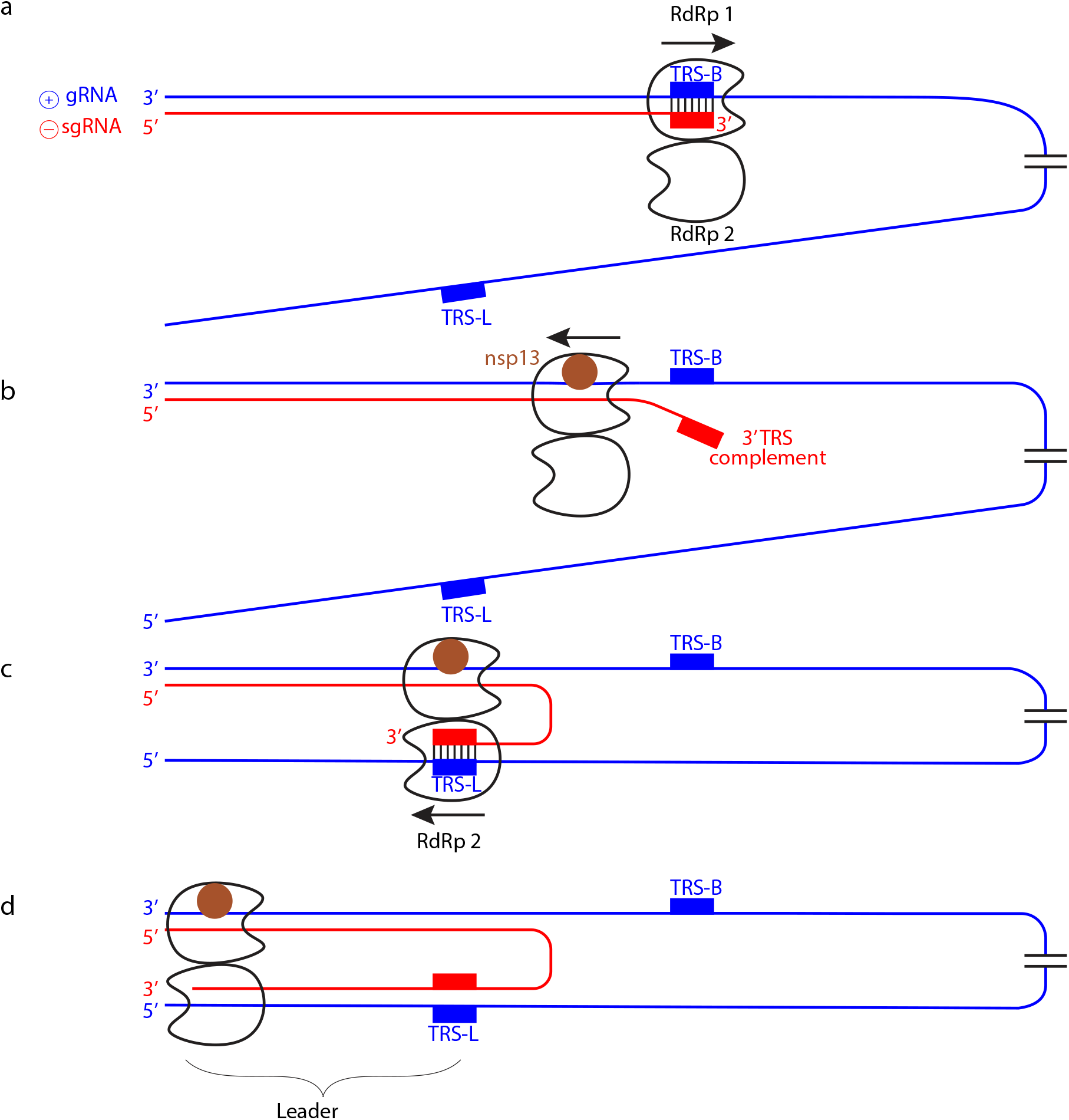
Model of subgenomic RNA production for transcription. **a** Genomic positive-strand (⊕) RNA (gRNA) is used as template to produce the 5’ region of negative-strand (⊖) sgRNA until TRS-B is reached by the RdRp monomer 1. **b** Backtracking is mediated by nsp13 helicase and exposes the newly synthesized, complementary TRS sequence. **c** The complementary sequence in sgRNA can pair with the downstream TRS-L in gRNA and is then loaded into RdRp monomer 2. **d** RdRp 2 then completes ⊖ sgRNA synthesis while RdRp 1 backtracks further.

In our model, it is not the RdRp that switches to a second RNA position, but instead the RNA switches to a second RdRp. After the switch, RdRp 1 may backtrack further until it reaches the 3’-end of the template, whereas RdRp 2 could move forward until it reaches the 5’-end of the template. These movements occur on the same template but in opposite directions and would be facilitated by the antiparallel arrangement of the polymerases. In support of such opposite movement, only one copy of the template-strand engaged nsp13^33^ can be modeled on our dimer structure, and thus only one RdRp in the dimer is predicted to backtrack (**Supplemental Figure 2c**).

## Methods

### Initial detection of RdRp dimers in cryo-EM data

To analyze the statistical distribution of RdRp monomers in our cryo-EM data, we calculated the nearest-neighbor (NN) distances and the relative orientations for all neighboring RdRp complexes using the previously refined monomer poses. We chose to express relative orientation through a single angle by calculating the angle of the rotation around the eigenvector of the rotation matrix. This showed that certain distances and relative orientations between two monomers were highly prevalent (**Figure 1**) and indicated the presence of dimeric particles where the two RdRps would adopt a specific distance and relative orientation with respect to each other. We then located such RdRp dimers in micrographs by identifying pairs of RdRps within a narrow range of NN distances and orientations. We observed the same overrepresented RdRp distance of 80 Å and relative angle of 180°, indicating the occurrence of a defined RdRp dimer. Furthermore, we could observe that filtering for relative orientations alone already enriched for the most abundant relative distance, further supporting the presence of an ordered dimer (**Supplementary Figure 1b**). We initially obtained ~31,000 dimeric particles using a distance <90 Å and relative orientation larger than 166° as a filtering criterion.

### Detection of additional dimeric particles

Because the yield of dimeric particles depended on both halves of a dimer being first detected as monomer, we aimed to detect more RdRp monomers using two strategies. First, we carried out template-based picking in RELION^34^ using the monomeric RdRp-RNA structure^10^ filtered to a resolution of 30 Å as a 3D reference to pick further monomers that might have been missed due to their proximity to other particles. Template-based picking of monomeric RdRp does however not introduce any model bias that could influence the dimer structure. Second, we used the previously established NN distance and relative orientation to predict for each RdRp monomer the position where its partner in a dimeric particle should be located on the micrograph. From our NN search we obtained the 3D offset that the second monomer should adopt in a dimeric particle (**Supplementary Figure 1b**) and used this to predict the monomer poses from our first refinement. This approach does not bias the analysis towards a fixed relative orientation of NN monomers. Instead, we calculated the orientation of each monomer *de novo* after combining all picked monomers from Warp, template-based picking and our prediction approach. After removing duplicates, we used the 3D classification approach described previously^34^ and conducted 3D refinement. Using the new monomer populations and their refined poses, we could predict 78,787 dimer particles.

### RdRp dimer reconstructions and model building

Final dimer coordinates were used to extract particles using a box size of 300 Å and a pixel size of 1.201 Å. 2D classification in cryoSPARC^35^ (**Supplementary Figure 2c**) showed a dimeric structure that contained two RdRps in antiparallel orientation. We selected 2D classes that showed two RdRp-RNA monomers without a gap between them. After 2D classification, we used 27,473 particles for an *ab initio* refinement with three classes. For the subsequent 3D refinement, we chose a reconstruction from the *ab initio* refinement that had two clearly resolved RdRp monomers close to one another as a reference. A molecular model was obtained by placing two copies of RdRp-RNA complexes (PDB-6YYT)^10^, removal of mobile regions and fitting of domains as rigid bodies in UCSF Chimera^36^.

## Data availability

Maps and models for the SARS-CoV-2 RdRp dimer structure have been deposited in the EM data base and Protein Data Bank (PDB) under accession codes XXXX and YYYY, respectively.

## Acknowledgements

P.C. was supported by the Deutsche Forschungsgemeinschaft (EXC 2067/1 39072994, SFB860, SPP2191) and the ERC Advanced Investigator Grant CHROMATRANS (grant agreement No 882357). H.S.H. was supported by the Deutsche Forschungsgemeinschaft (EXC 2067/1 39072994, SFB1190, FOR2848).

## Contributions

FAJ carried out data analysis, assisted by DT. HSH and GK helped with modeling. HSH, GK, JS and CD helped with structure interpretation, PC supervised the project and wrote the manuscript, with input from all authors.

---

**Supplementary Figure 1.**
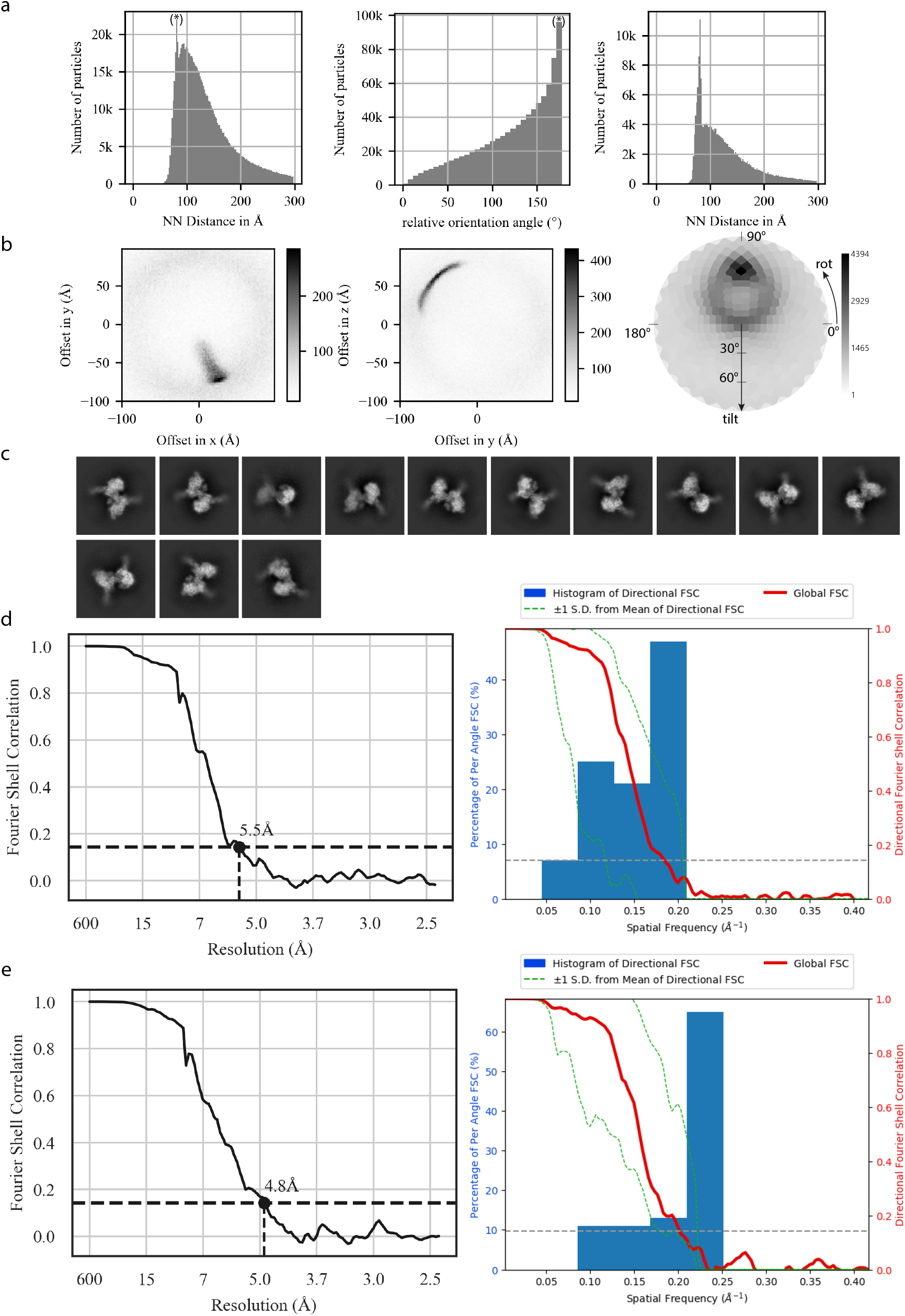
Determination of dimeric RdRp structure. **a** Detection of RdRp dimers in cryo-EM data. On the left, nearest neigbor (NN) distances for all Warp-picked monomeric RdRp particles, with the most significant peak that deviates from the underlying random distribution indicated (*) at around 75 Å. In the middle plot, NN orientations are shown, expressed as a single angle of rotation around the eigenvector of the rotation matrix. A clear peak (*) is visible close to 180°. On the right, a histogram of NN monomer distances, where only those nearest neighbors that also have a relative orientation of >166° are considered. This sharpens the nearest neighbor distance peak, indicating a correlation between this distance and a defined orientation within dimers. **b** Detailed analysis of our peak distances and relative orientations. In the first two panels, the elements of the vector connecting two NN monomers, expressed in the reference frame of one of the two monomers, are shown, further indicating a defined co-localization of two monomeric RdRps. In the third panel, a projection of relative orientations on a half sphere are shown. **c** Selected 2D class averages from our predicted 78,787 dimeric particles. We chose classes with two clearly defined RdRp monomers and strong RNA signal as an indication for high alignment quality, representing 27,473 particles in total. **d** Fourier shell correlation (FSC) curve for the masked reconstruction of the antiparallel dimer, indicating an average resolution of 5.5 Å. The second panel contains a histogram of directional FSC values as calculated by the 3DFSC server^37^, indicating significant anisotropy. **e** With C2 symmetry applied, the average resolution of the reconstruction increased to 4.8 Å. The second panel contains a histogram of directional FSC values as calculated by the 3DFSC server^37^, indicating significant anisotropy.

**Supplementary Figure 2.**
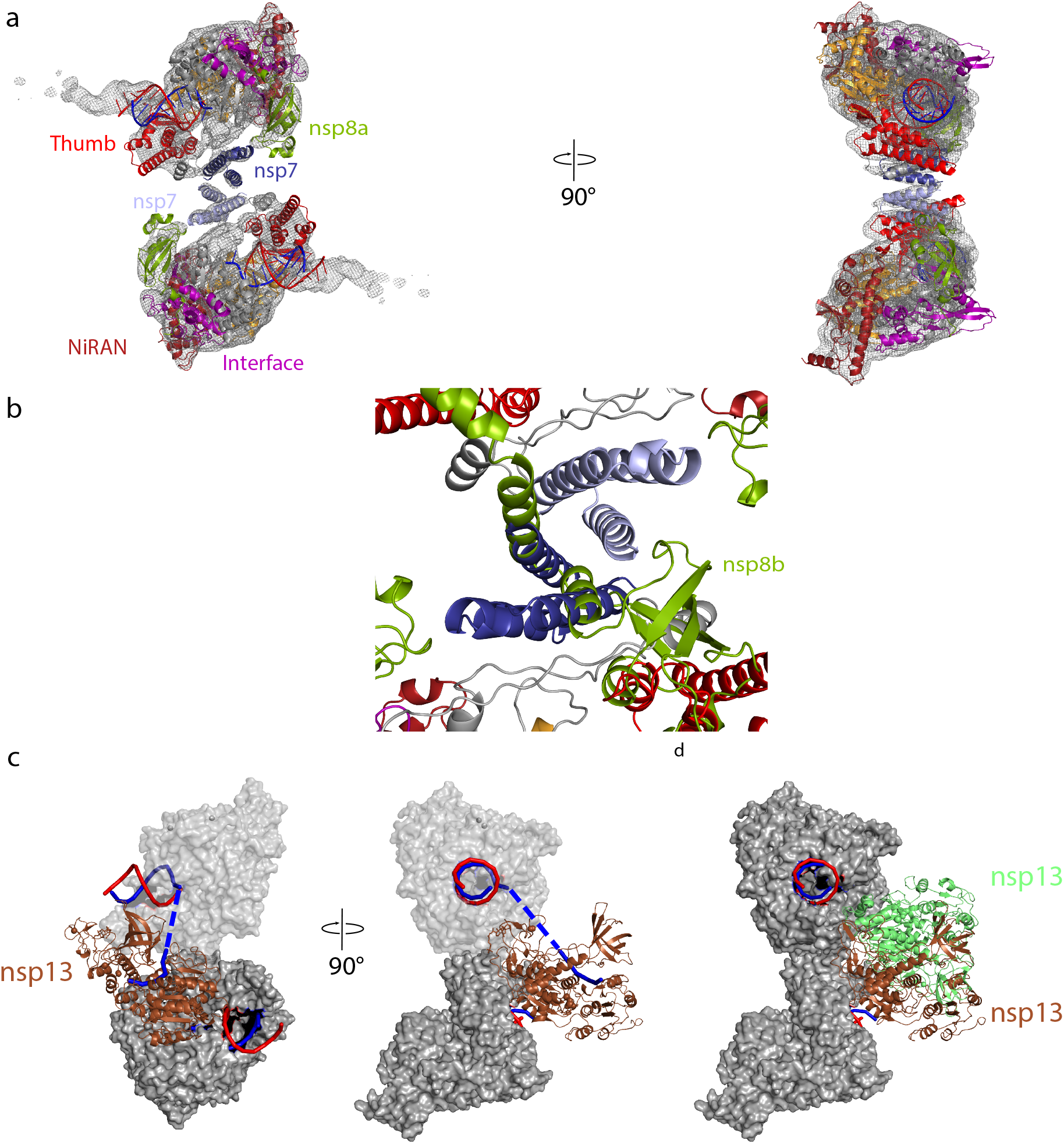
Quality of dimeric RdRp structure and structural comparisons. **a** Fit of structural model to cryo-EM density. Density was weak or lacking for the second turn of RNA and the associated nsp8a sliding pole. Views are related by a 90° rotation around the vertical axis. **b** Modeling the second nsp8 copy (nsp8b) onto the top RdRp in the dimer structure resulted in clashes with the nsp7 subunit of the neighboring RdRp on the bottom. **c** Modeling shows the nsp13 helicase may be accommodated on the RdRp dimeric structure. The nsp13 copy nsp13.1 (chain E in 7CXN^33^) can bind to one of the two monomers (binding to top monomer shown). Possible trajectory of 5’ template RNA is indicated in the second panel as a dashed blue line. **d** nsp13.1 cannot bind to both monomers. Modelling one copy bound to the top monomer (brown) and one bound to the bottom monomer (lime) shows clashes between them.

